# AstraROLE2 & AstraSUIT2: Multi-Task Annotation Models for Functional Profiling of Proteins

**DOI:** 10.1101/2025.06.21.660734

**Authors:** Çağlar Bozkurt, Alexandra Vasilyeva, Aniruddh Goteti

## Abstract

Most *in-silico* protein characterisation tools focus on only one aspect of protein function, forcing researchers to use multiple models or to bypass computational checks. Here we introduce AstraROLE2 and AstraSUIT2, two transformer-based, multi-task annotators that deliver an integrated functional profile in a single pass.

A 1,351-dimensional input (ESM-2 CLS embeddings plus physicochemical Orbion enrichments) is mapped by a 512-unit encoder and task-specific linear heads: four in AstraROLE2 (EC class, GO term, molecular pathway, protein category) and nine in AstraSUIT2 (cofactor group, specific cofactor, domain, host, membrane association type, transmembrane helix number, subcellular localization, quaternary category, quaternary stoichiometry). Models were trained on 730k UniProt proteins with stratified 70/15/15 splits; class-weighted BCE and Optuna hyper-parameter search countered imbalance. On hold-out sets the heads reached macro F_1_=0.84–0.98 and MCC=0.85–0.98. Highest scores were seen for cofactor binding (0.98), membrane association type (F_1_=0.97) and top-level EC number (0.96); GO term classification was hardest (0.85). Against recent comparators (incl. DeepGOPlus and TargetP 2.0), the Astra models matched or exceeded performance, especially on metal-ion binding and cofactor binding. Additional tests on three novel proteins not included in initial dataset showed good predictions for most labels, underscoring the potential for hypothesis generation.

Overall, AstraROLE2 and AstraSUIT2 supplied fast, state-of-the-art multi-label protein annotation within one unified model network.

## 1 Introduction

Using AI-driven insights for biomedical experiments and protein design is rapidly gaining traction. Artificial intelligence is leveraged to accelerate drug discovery, identify novel biomarkers, process microscopy and scattering data, and facilitate the automation of laboratory workflows.(1) In light of the long time-scales and high failure rates that characterise the drug discovery pipeline, it is hoped that utilising AI can improve both the speed and the success rate of developing novel drug candidates and studying potential drug targets.(2; 3)

A number of tools are available to researchers that can assist with stabilising proteins and interpreting experimental outcomes.(1; 2; 3) However, many of the currently available models focus on only one aspect of protein biology, such as post-translational modifications (PTMs) or taxonomic-classification prediction, and integrating insights from a large number of models can be challenging. At the same time, multiple protein characteristics need to be considered: for instance, taxonomy and PTMs influence the choice of expression system, whereas the expected molecular function informs the necessary assays.

If modifications to the protein are introduced, it is desirable to assess all its parameters *in silico* before committing to experimental work; within a single structure-determination project, dozens of mutant variants may be screened for stability, making the use of multiple input models impractical or impossible. As a result, mutant proteins are often created via rational design and tested experimentally without a reliable intermediate assessment, introducing delays and extra costs into the process in case of failure.

Our goal is to create a family of Orbion AI models using state-of-the-art methods and to integrate them seamlessly within a single platform, thereby providing an accurate, multifaceted overview of protein features. Previously, we introduced the AstraPTM model, which employed a transformer-based architecture to predict PTMs with high reliability; the AstraPTM2 model is currently being prepared for publication. Here, we report on two additional AI models—AstraROLE2 and AstraSUIT2—that aim to provide a more rounded picture of a protein by predicting its likely biological role along with key features such as metabolic-pathway membership and sub-cellular localisation.

## 2 Methods

### Architecture

AstraROLE2 and AstraSUIT2 are both single-encoder, multi-task **transformers**(9) that turn the joint protein feature vector (protein-level embedding + ESM2 CLS embeddings) into functional annotations in one forward pass (Figure 1). Briefly, the input features are projected to a Linear Layer, sent to a Positional Encoding layer, and then sent through the Transformer encoder layers, which are connected to different heads for each task, with sigmoid functions. AstraROLE2 and AstraSUIT2 represent updated versions of previously reported AstraROLE and AstraSUIT, with the same overall architecture and expanded label sets. The step-by-step architecture of the models is described in more detail in the Appendix.

**Figure 1:**
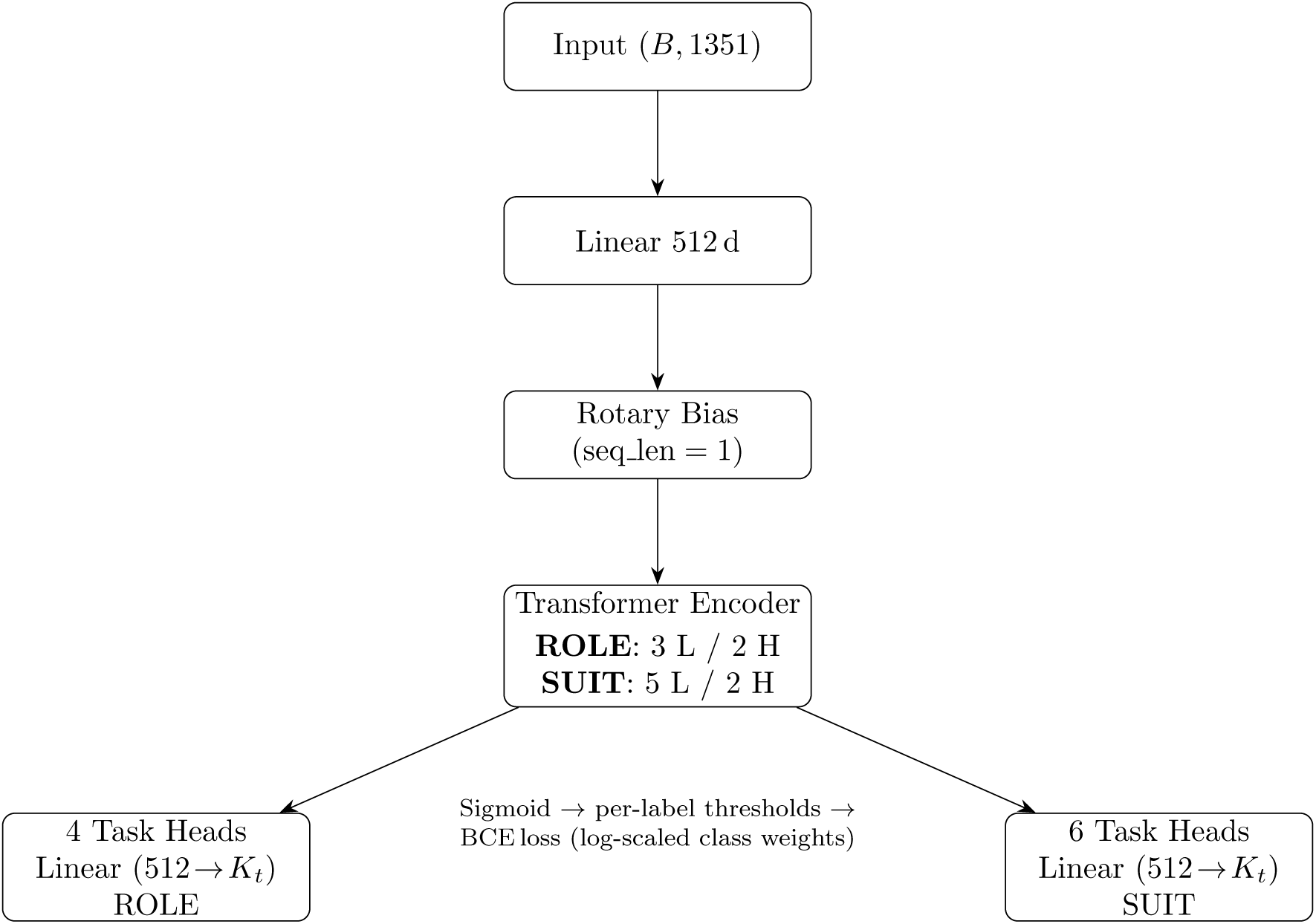
Vertical data-flow diagram for the Astra backbone and its two model-specific task-head branches. L = transformer layers, H = attention heads.

### Training

Both models accept as input a 1,351 dimensional vector per protein, including protein sequence, ESM2 embeddings and Orbion enrichments. The dataset comprised 730,197 unique proteins curated from **UniProt**(8) (Swiss- and TrEMBL; database cut-off 31.12.2024; see Figure 2 for top-line dataset overview and Appendix for more detailed classification).

**Figure 2:**
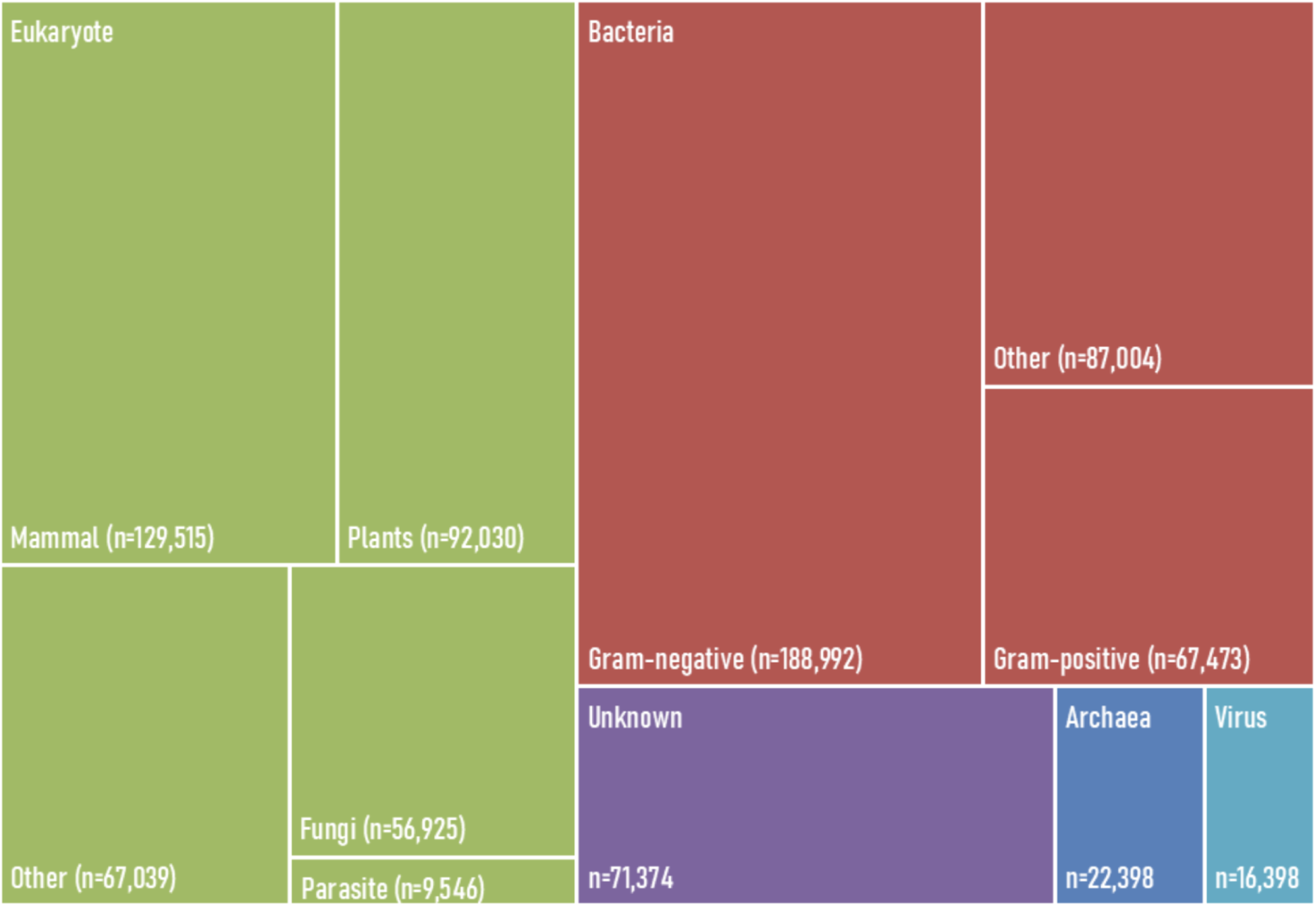
Overview of protein taxonomy labels in the UniProt dataset.

Data were retrieved and categorized via UniProt’s API into individual JSON files; 12,342 isoforms were not included due to their data inconsistencies from the UniProt JSON API. Then, through a proprietary classifier, comments, annotations, categorized information, references, and keywords were parsed to find the right values for each label.

ESM2 embeddings, which help to incorporate evolutionary information, were generated from esm2 t33 650M UR50D(6). CLS versions were used here, as token representations[0]. To add descriptive information on the protein, the enrichments from Orbion were added that include molecular weight, GRAVY score, isoelectric point, length and number of bonds.

Normalization was performed through a StandardScaler on the training data. Both mean and standard deviation values were saved into scaler files, which were then used on inference requests on the models. The data set was shuffled and split 70% for training, 15% for validation, and 15% for testing, independently for each model.

Multiple training iterations were run for each model to optimize the performance of the predictions. On the protein feature set approaches, CLS, LSE, and Mean ESM embedding versions were tested, where the F_1_ scores for all heads were checked. As these tests didn’t show a significant difference between different features, CLS was used as the key ESM feature set approach. After this, hyper-parameter optimisation was made using **Optuna**(4).

## 3 Results

### Overall Performance

Both AstraROLE2 and AstraSUIT2 models showed good overall performance, with macro F_1_ for label groups between 0.853 and 0.982 and macro Matthews Correlation Coefficient (MCC) between 0.851 and 0.977 (see Table 1). The most accurate category for prediction was cofactor binding (F_1_=0.982, MCC=0.977), followed closely by membrane localisation (F_1_=0.967, MCC=0.960) and top-level EC number (F_1_=0.958, MCC=0.953). Lowest F_1_ was seen for GO terms (F_1_=0.853, MCC=0.852).

**Table 1:** Overall performance of AstraROLE2 and AstraSUIT2 models.

| Model | Output | Acc | Prec | Rec | F<sub>1</sub> | MCC |
| --- | --- | --- | --- | --- | --- | --- |
| ASTRAROLE2 | EC Number | 0.991 | 0.961 | 0.956 | 0.958 | 0.953 |
|  | GO Terms | 0.996 | 0.874 | 0.834 | 0.853 | 0.852 |
|  | Protein Categories | 0.969 | 0.875 | 0.861 | 0.868 | 0.851 |
|  | Pathway Memberships | 0.997 | 0.945 | 0.898 | 0.921 | 0.92 |
| ASTRASUIT2 | Cofactor Binding | 0.993 | 0.982 | 0.982 | 0.982 | 0.977 |
|  | Cofactor Specific | 0.999 | 0.95 | 0.921 | 0.936 | 0.935 |
|  | Domain Classification | 0.981 | 0.866 | 0.869 | 0.868 | 0.858 |
|  | Organism Association | 0.974 | 0.869 | 0.906 | 0.887 | 0.873 |
|  | Membrane Type | 0.989 | 0.965 | 0.968 | 0.967 | 0.96 |
|  | Subcellular Location | 0.984 | 0.882 | 0.864 | 0.873 | 0.864 |
|  | TM Helices Group | 0.994 | 0.877 | 0.878 | 0.878 | 0.875 |
|  | Quaternary Category | 0.973 | 0.886 | 0.856 | 0.871 | 0.856 |
|  | Quaternary Stoichiometry | 0.993 | 0.917 | 0.875 | 0.895 | 0.892 |

ACC: Accuracy, Prec: Precision, Rec: Recall; MCC: Matthews Correlation Coefficient. Green = Best, Yellow = Moderate, Red = Weakest (within one column)

### Per-Label Performance

Performance metrics (accuracy, precision, recall, F_1_, MCC) for each label are provided in the Appendix. Here we describe the key outcomes, focusing on F_1_ as a representative metric due to the imbalance of the dataset.

For top-level EC number, F_1_ varied from 0.987 for ligases to 0.933 for hydrolases, with much lower F_1_=0.653 for catch-all term “Unclassified enzyme”. In protein category prediction, enzymes (F_1_=0.968) and transporters (F_1_=0.928) performed best, and the worst performance was in signaling and storage proteins (F_1_=0.697 for each).

Prediction for pathways was overall accurate, with lower performance in sphingolipid metabolism (F_1_=0.695) and the mevalonate pathway (F_1_=0.747). Interestingly, the best model performance was seen for the non-mevalonate (MEP) pathway (F_1_=0.983), which is an alternative to the mevalonate pathway found in bacteria and plants, as well as for the Shikimate pathway (F_1_=0.983).

In GO term prediction, six labels did not return relevant hits and therefore returned null F_1_; among other labels, lowest performance was seen for translation factor activity, cell adhesion molecule binding and biotin binding (F_1_*≤* 0.400). For other labels, performance was satisfactory, with highest F_1_ for ATP binding (F_1_=0.968) and GTP binding (F_1_=0.966; see Figure 3).

**Figure 3:**
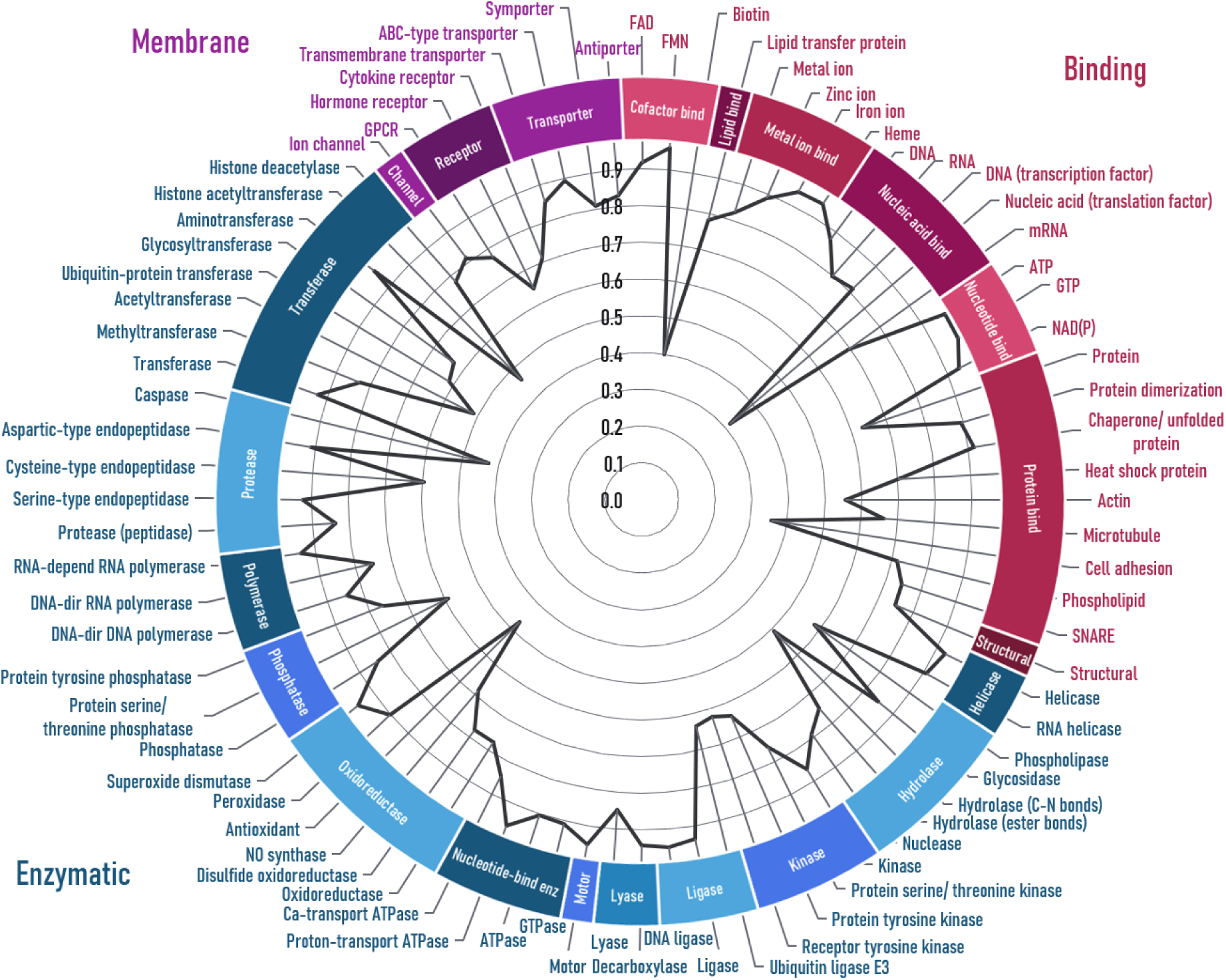
F_1_ factor for AstraROLE2 performance per GO term. Six labels with null F_1_ are not shown.

In AstraSUIT2, cofactor binding was assessed on two levels: general (e.g. organic, metal ion) and specific (e.g. Ca, Fe, biotin, heme). In general cofactor prediction, F_1_*>*0.9 was seen for all labels. For specific cofactors, seven labels returned null results, and other labels varied from F_1_=0.300 (pyrroloquinoline quinone) to F_1_=1.000 (guanosine monophosphate; see Figure 4).

**Figure 4:**
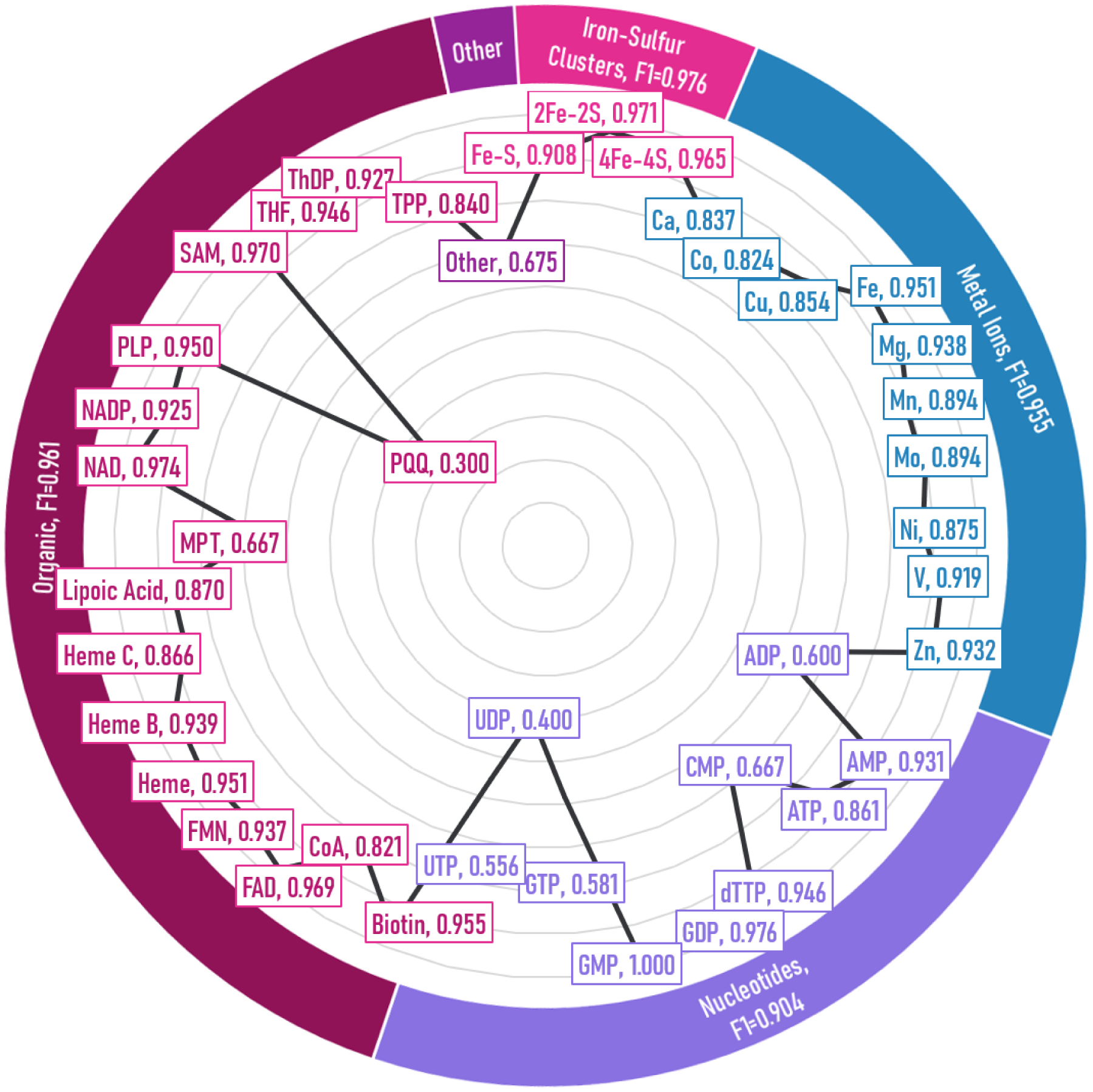
F_1_ factor for AstraSUIT2 performance per cofactor label. Labels with null F_1_ are not shown. ADP: Adenosine Diphosphate, AMP: Adenosine Monophosphate, ATP: Adenosine Triphosphate, CMP: Cytidine Monophosphate, CoA: Coenzyme A, dTTP: Deoxythymidine Triphosphate, FAD: Flavin Adenine Dinucleotide, FMN: Flavin Mononucleotide, GDP: Guanosine Diphosphate, GMP: Guanosine Monophosphate, GTP: Guanosine Triphosphate, MPT: Molybdopterin, NAD(P): Nicotinamide Adenine Dinucleotide (Phosphate), PQQ: Pyrroloquinoline Quinone, PLP: Pyridoxal Phosphate, SAM: S-Adenosylmethionine, ThDP: Thiamine Diphosphate, TPP: Thiamine Pyrophosphate, THF: Tetrahydrofolate, UDP: Uridine Diphosphate, UTP: Uridine Triphosphate

Associated organisms were classified in two groups of labels: domain subgroups (e.g. gram-positive bacteria) and organism/ host (e.g. animal pathogen, animal-associated). Domain classification returned null F_1_ for the synthetic protein category and overall lower F_1_ for catch-all “other” labels; best performance was for gram-negative bacteria (F_1_=0.938). Organism/ host F_1_ was from 0.347 (bacterial pathogen) to 0.971 (bacteriaassociated).

Membrane association type was predicted reliably (F_1_=0.824–0.983), aside from the lipid-anchored and GPI-anchored labels, which returned null results. Number of transmembrane helixes was also estimated well (F_1_=0.810–0.925). Membrane labels also showed best performance in the subcellular localisation category (F_1_=932 for membrane, F_1_=0.921 for inner membrane), and all other subcellular labels performed well with F_1_*>*0.75 (see Figure 5).

**Figure 5:**
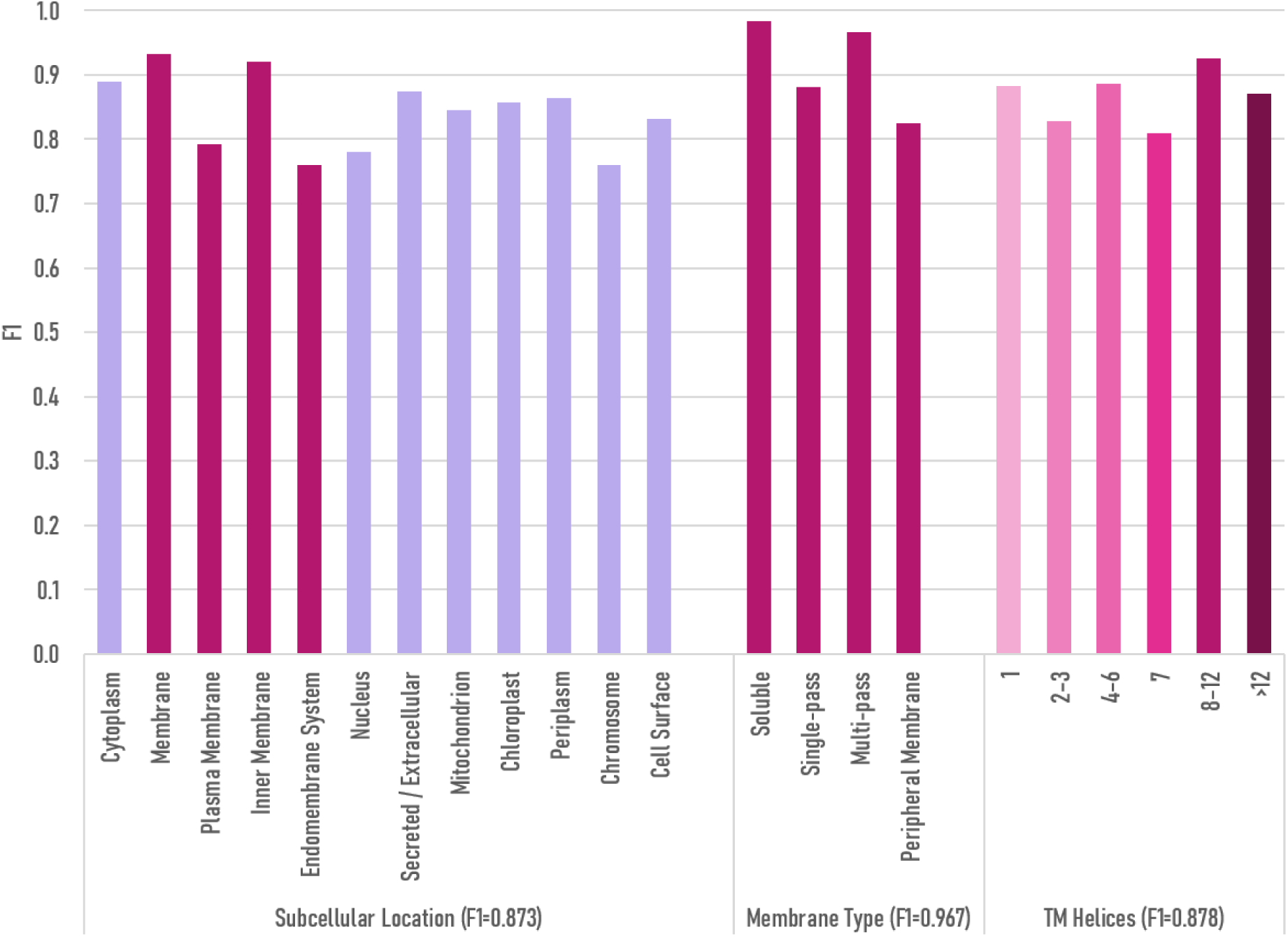
F_1_ prediction metrics for subcellular location and membrane labels.

Finally, two groups of labels assessed quaternary structure: one for general categorisation and one for the predicted number of protein units (stoichiometry). For all labels in these groups prediction was reliable with F_1_*>*0.8.

### Case Studies

In order to additionally assess the performance of the models, we identified three proteins which were not included in the UniProt database before the download date (31.12.2024) and that were reported to have a high degree of structural and/or functional novelty.

IgtG is a glycosyltransferase from gram-positive bacteria that catalyzes iterative addition of up to four glucose units to a single serine on mature lasso peptides.(10) IgtG and its homologs are evolutionally separate from other known ribosomally synthesized and post-translationally modified peptide (RiPP) glycosyltransferases. The Astra models correctly identified the bacterial taxonomy but suggested sub-classification as “other” (90% certainty). Membrane association-related labels were predicted with almost 100% certainty, indicating a 4-6 transmembrane helix protein, which aligns with structural predictions. Transferase function was identified correctly both as a top-level EC class (73%) and GO term (76%). Although the specific label for peptidoglycan biosynthesis did not reach significance, higher-level enzyme and structural functions were identified with a high level of confidence (*>*90%).

esmGFP is a synthetic fluorescent protein that shares 58% sequence identity with the closest known natural fluorescent protein.(11) The models did not return a positive taxonomy but suggested animal interaction (31%), which aligns with a novel synthetic GFP variant. Other predictions were soluble (82%), located and a specialized function (69%), with none of the evaluated GO terms or molecular pathways returning a positive hit. All of these align with expectation for a synthetic fluorescent protein and show no false positives.

TPM3P9 is a human microprotein previously believed to be associated with noncoding RNA; it regulates RNA splicing with a potentially oncogenic effect and lacks significant sequence similarity to known proteins.(12) It was identified by the Astra models as a human-associated (88%) soluble (99%) mammalian protein (91%). The model provided no false positives on EC and GO terms, but did not identify the regulatory function (predicting instead structural and specialised/ other function).

## 4 Comparisons

AstraROLE2 and AstraSUIT2 were benchmarked against the newest publicly-available protein-annotation models released between 2022–2025, where a comparison was possible. The following models were included as potentially relevant comparators:

- **GO-Slim** : TALE+(13), DeepGO-SE(14), GNN-GO(15), ProteInfer(21)
- **EC superclass** : GraphEC(16), DeepEC-Transformer(17), ECPred-GAT(18)
- **Cofactor / metal binding** : M-Ionic(19), bindEmbed21(20), DeepCoFi(22)
- **Localisation** : DeepLoc 2.0(23), LightAttention-db10(24), DeepLoc-Pro(25), SCLpred-ECL(26)
- **Membrane / topology** : DeepLoc 2.1(27), DRHGNN(28), TMbed(29), DeepTMHMM(30)

The heterogeneity of comparator models makes direct comparisons challenging, in particular with regard to different labels and reported metrics. For GO terms, GO-Slim metrics were matched; for localisation, we used the 10- and 9-class label maps published with DeepLoc 2.x(23). Binding tasks were collapsed to *protein-level* labels to align residue-level baselines such as M-Ionic and bindEmbed21 with AstraSUIT2(19; 20). For result assessment, metrics were used as reported by comparator model publications (*F*_max_, macro- or micro-*F*_1_, segment-level *F*_1_, or per-protein accuracy), and the Astra value that corresponds most closely was provided in each case.

## 5 Discussion

The model performance was overall satisfactory, with some areas such as top-level EC class and membrane type labels showing particularly strong predictions. Several labels, such as “Bacterial pathogen” and “PQQ cofactor”, yielded low performance metrics compared with other labels in similar categories, likely due to low count numbers; training data will be enhanced in future model iterations. The performance on the other labels has shown that, with a larger sample set on these labels, the performance can further improve.

Direct comparison of AstraSUIT2 and AstraROLE2 with other models was not always feasible due to the broad heterogeneity in output labels and performance metrics that other models report. In addition, factors such as data split methods and embeddings had to be considered when interpreting result comparisons: Astra results are on a 70*/*15*/*15 random hold-out, which is typically 5–10pp easier than the strict H or temporal splits used by several competitors(13; 16). Nonetheless, it can be generalised that the last several years have seen a sharp improvement in model results, and that there is a strong tendency towards using residue-level embeddings, such as ESM-2(6), and the **Transformer** architecture(9); this validates the architecture approach chosen for the Astra models.

Where direct comparison was feasible for some of the output metrics, Astra models performed comparably or often superior to comparator models. Top-level metrics for prediction of metal ion binding were particularly strong in the Astra models versus two comparators (see Table 2). Although these comparisons are by necessity limited by differences in evaluated labels and metrics, this nevertheless points towards strong performance by AstraROLE2 and AstraSUIT2.

**Table 2:** Astra versus recent specialist models. Split codes: R = random, H = homology, <u>B = balanced. Per-row best scores in bold. All scores are</u> higher = better.

| Task & Metric | Model | Astra | Split | Compared Model | Score |
| --- | --- | --- | --- | --- | --- |
| GO ($F_{\max}$) | ROLE2 | **0.97** | H | TALE+ (2023) | 0.78 |
| EC Superclass (Macro-$F_1$) | ROLE2 | **0.96** | R | GraphEC (2024) | 0.95 |
| Metal-ion Binding ($F_1$) | SUIT2 | **0.96** | H | M-Ionic (2024)<sup>†</sup> | 0.53 |
| All Cofactors (Macro-$F_1$) | SUIT2 | **0.98** | R | DeepCoFi (2024) | 0.74 |
| Membrane Type (Micro-$F_1$) | SUIT2 | **0.98** | H | DRHGNN (2023) | 0.89 |
| TM-Helix Group (Seg. $F_1$) | SUIT2 | 0.87 | H | TMbed (2023) | **0.89** |
| TM-Helix Count (Accuracy) | SUIT2 | **0.99** | H | DeepTMHMM (2022) | 0.93 |

<sup>†</sup>Residue-level metric; AstraSUIT value is protein-level, numbers are therefore only indicative.

Three case studies were performed on particularly challenging proteins that were not included in the UniProt dataset and were reported to have a high degree of novelty (IgtG, esmGFP and TPM3P9). The models yielded false negatives on some labels, but for most labels the performance was good, especially considering protein novelty; IgtG was correctly predicted to be a multi-pass membrane protein with transferase activity; esmGFP predictions fully aligned with a new synthetic protein analogous to GFP; and TPM3P9 was classified as a human-associated soluble protein with high confidence. These results highlight the potential for Astra models to assist in hypothesis generation (for example in exploring the dark genome) and in performing checks on designed mutated constructs and de novo proteins.

Next steps in developing the Astra models will involve incorporating more training data, both by volume and type, for example examining whether the addition of structures to the input would substantially improve outcomes. We will also continue investigating the potential for synergy between different models to provide better and more informative predictions of protein behaviour.

## 6 Model Access

AstraROLE2 and AstraSUIT2 can be reached through two complementary entry-points:

- **Interactive Demo — Free, Rate-limited** The models are available at https://app.orbion.life/astra/. No registration is required for rate-limited testing.
- **Research API — Unlimited Throughput** For large-scale or automated studies, an account is required at https://apporbion.life/ with free academic access and license options for commercial use.

## 7 Conclusions

AstraROLE2 and AstraSUIT2 models showed good overall performance, with F_1_=0.853–0.982 for label group macro-F_1_, and best predictions for membrane type, top-level EC number and cofactor binding. On label level, several label clusters showed particularly good predictions (F_1_*>*0.9), but there were also several outlier labels with low performance. Where comparison with other models was possible, AstraROLE2 and AstraSUIT2 showed mostly superior performance. These results support our choice of architecture and will inform the development of further models in the Astra family. Due to the good predictive performance of AstraROLE2 and AstraSUIT2, these models can be used to assist with hypothesis generation and to support the experimental research process.

## A Appendix

### UniProt Dataset

**Table 3:**
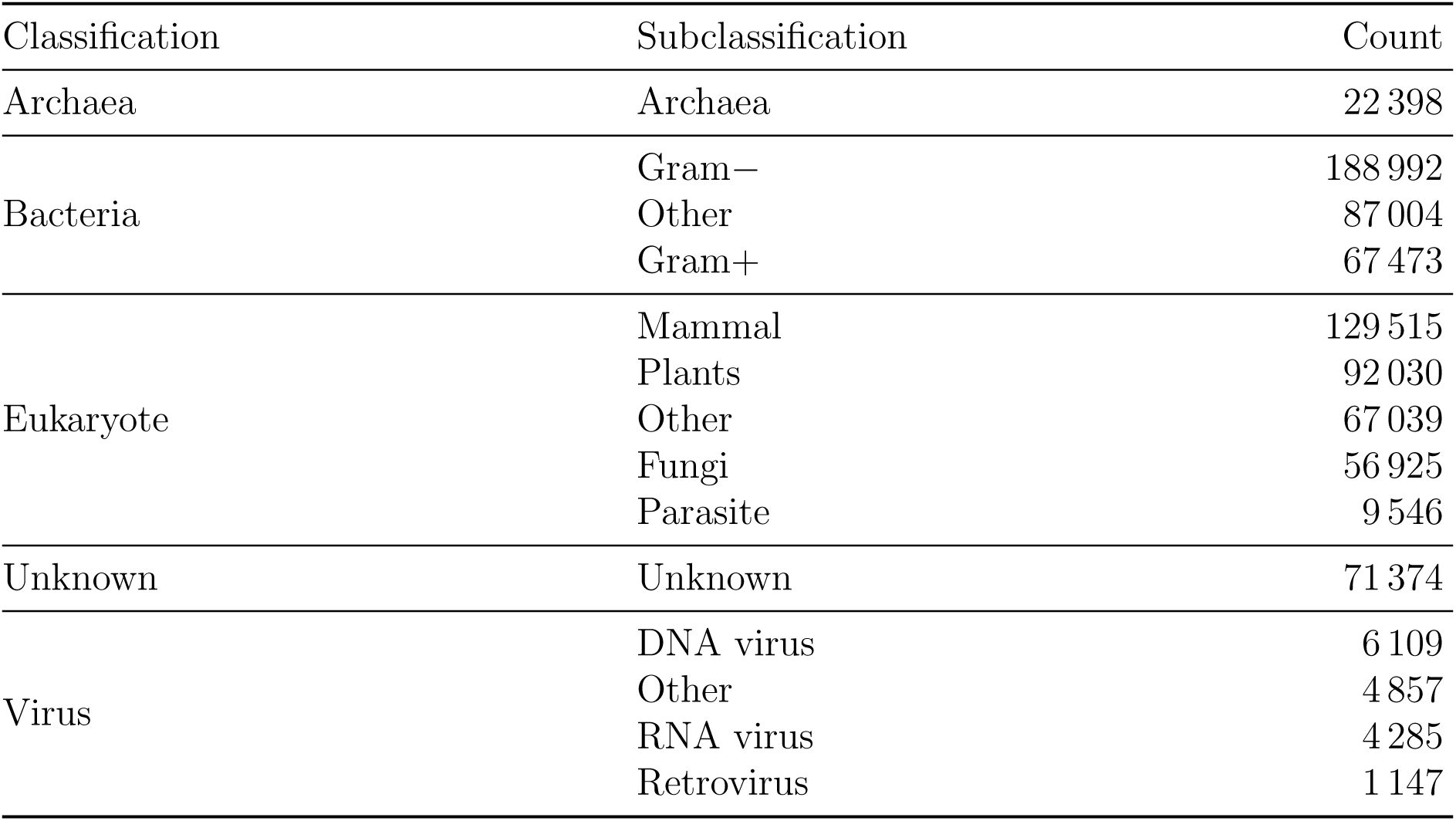
Taxonomic distribution of sequences.

**Table 4:** Functional categories and counts.

| Classification | Category | Count |
| --- | --- | --- |
| Cofactors | Unknown | 496 821 |
|  | Organic cofactors | 170 620 |
|  | Metal ions | 149 470 |
|  | Iron-Sulfur clusters | 14 556 |
|  | Others | 7 352 |
| Membrane type | Soluble | 666 750 |
|  | Multi-pass | 66 926 |
|  | Single-pass Type II | 34 940 |
|  | Lipid-anchored | 2 313 |
|  | GPI-anchored | 1 327 |
|  | Single-pass Type I | 705 |
|  | Peripheral membrane | 46 |
| Organism association | Animal | 526 393 |
|  | Unknown | 284 788 |
|  | Plant | 74 187 |
|  | Human | 62 378 |
|  | Insect | 16 523 |
|  | Environmental | 2 851 |
|  | Others | 2 239 |
|  | Commensal/Symbiont | 1 444 |

#### Architecture

**Table 5:** Exact layer configuration for the two Astra models. *B* = batch size, *D* = 1 351 input features, *K_t_* = task-specific output width.

| Stage | AstraROLE | AstraSUIT |
| --- | --- | --- |
| Input Tensor | $(B, D)$ | $(B, D)$ |
| Linear Projection | $(B, 1, D) \rightarrow (B, 1, 512)$ | $(B, 1, D) \rightarrow (B, 1, 512)$ |
| Positional Bias | Rotary (Sequence Length = 1) | Rotary (Sequence Length = 1) |
| Transformer Encoder | 3 Layers, 2 Heads, Model Dim 512 | 5 Layers, 2 Heads, Model Dim 512 |
| Shared Representation | $(B, 1, 512)$ | $(B, 1, 512)$ |
| Task Heads | $4 \times \text{Linear}(512 \rightarrow K_t)$ | $6 \times \text{Linear}(512 \rightarrow K_t)$ |
| Activation | Sigmoid | Sigmoid |
| Thresholding | Per-label Optimal Cut-off | Per-label Optimal Cut-off |
| Training Loss | BCE w/ Log-scaled Class Weights | BCE w/ Log-scaled Class Weights |

The architecture and optimisation for AstraROLE2 and AstraSUIT2 closely followed the same structure and metrics, with the only key difference being increased numbers of labels and label groups (i.e. heads).

#### Optimization and Hyperparameters

For AstraROLE, the optimization process led to 39 optimization trials on hyperparameters, where 22 of them were pruned in the process, and 16 were completed (see Table 1 for the selected hyperparameters). Overall, 40, 45, 50, and 60 epochs were tested; 50 epochs was the optimal point, where a higher number of epochs lead to an overfit. After 54 training iterations (incl. hyperparameter optimization), the model has shown a net growth of 0.10% on EC Number F_1_, 3.80% on GO Terms F_1_, 1.10% on Protein Categories F_1_, and 2.10% on Pathway Memberships F_1_.

For AstraSUIT, the optimization led to 26 trials on hyperparameters, with 7 pruned in the process and 19 completed. 40 and 60 epochs were tested and 40 epochs was the optimal point. After 41 training iterations (incl. hyperparameter optimization), the model has shown a net growth of 0.40% on Cofactor Binding F_1_, 2.40% on Domain-level Classification F_1_, 0.40% on Host Association F_1_, 1.80% on Subcellular Location F_1_, and 1% on TM Helices F_1_.

**Table 6:** Key hyper-parameters chosen for AstraROLE and AstraSUIT, optimised with Optuna.

| Hyper-parameter | AstraROLE | AstraSUIT |
| --- | --- | --- |
| Hidden dimension | 512 | 512 |
| Number of attention heads | 2 | 2 |
| Number of layers | 3 | 5 |
| Dropout rate | 0.10 | 0.21 |
| Training epochs | 50 | 40 |
| Learning rate | $5.96 \times 10^{-5}$ | $8.13 \times 10^{-5}$ |
| Weight decay | $9.16 \times 10^{-6}$ | $1.05 \times 10^{-5}$ |
| Loss-scale factor | 0.55 | 0.90 |
| Batch size | 64 | 128 |

**Figure 6:**
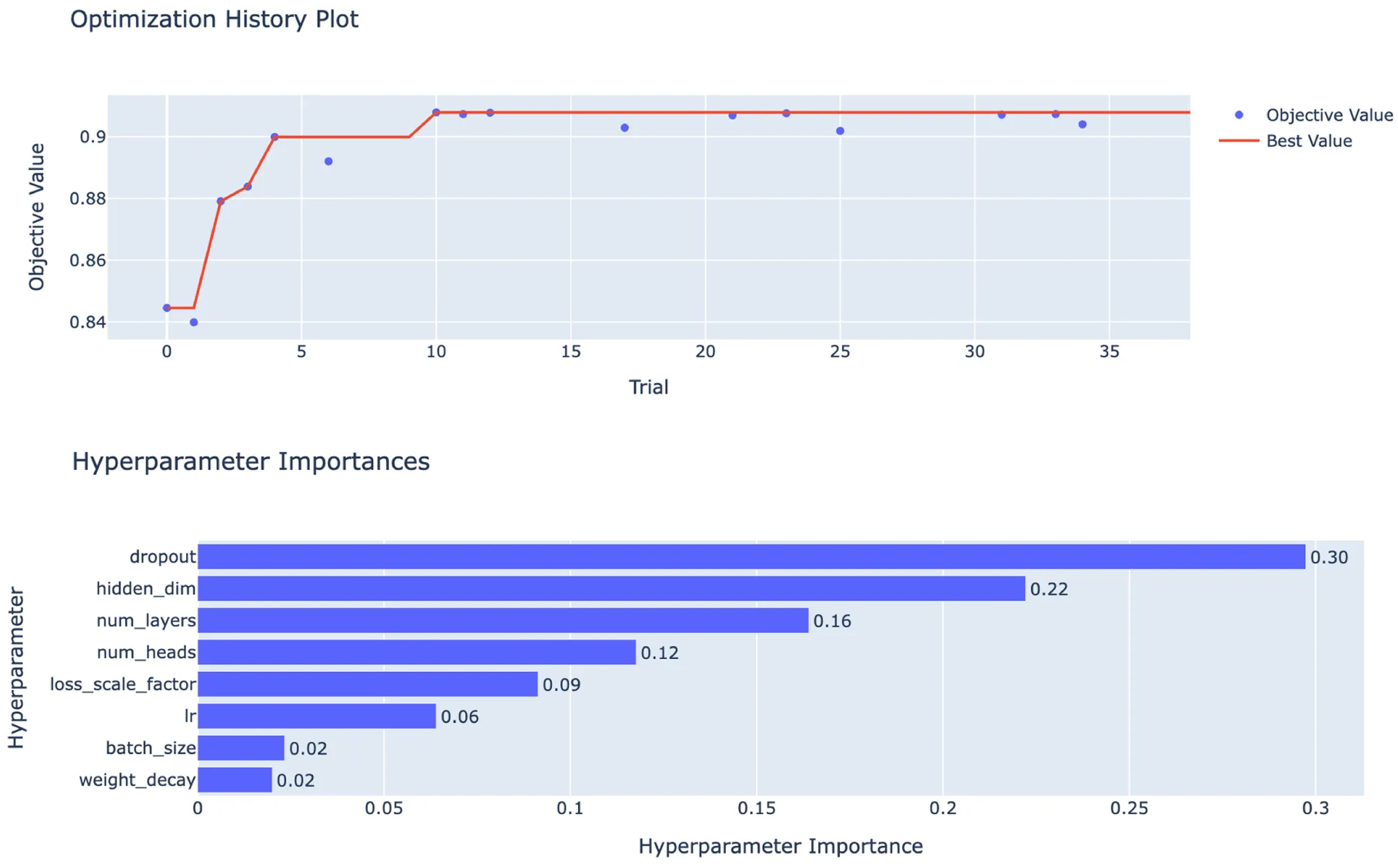
The historical performance and the hyperparameter importance rates based on the experiments ran by Optuna for AstraROLE.

**Figure 7:**
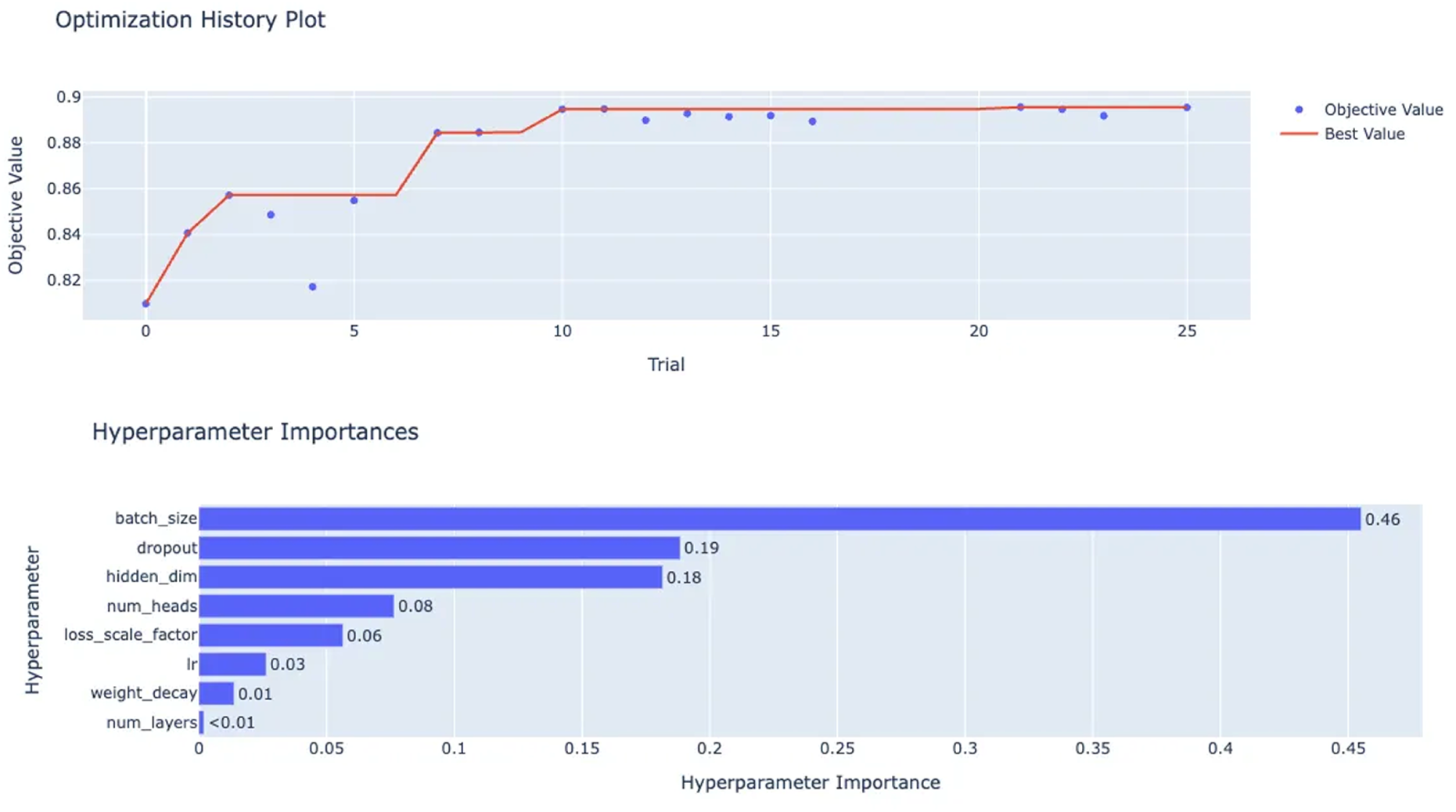
The historical performance and the hyperparameter importance rates based on the experiments ran by Optuna for AstraSUIT.

#### AstraROLE2 Results

**Table 7:**
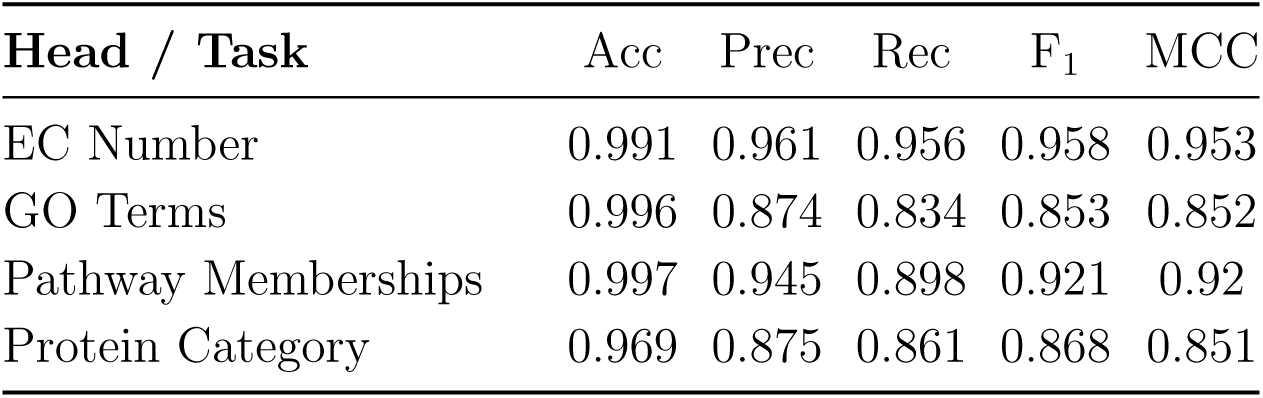
Performance of the four AstraROLE2 heads on the held-out test set.

| **Head / Task** | Acc | Prec | Rec | F<sub>1</sub> | MCC |
| --- | --- | --- | --- | --- | --- |
| EC Number | 0.991 | 0.961 | 0.956 | 0.958 | 0.953 |
| GO Terms | 0.996 | 0.874 | 0.834 | 0.853 | 0.852 |
| Pathway Memberships | 0.997 | 0.945 | 0.898 | 0.921 | 0.92 |
| Protein Category | 0.969 | 0.875 | 0.861 | 0.868 | 0.851 |

**Table 8:** Per-class performance of the EC-superclass head.

| **EC superclass** | Acc | Prec | Rec | F<sub>1</sub> | MCC |
| --- | --- | --- | --- | --- | --- |
| Oxidoreductases (EC 1) | 0.994 | 0.935 | 0.956 | 0.945 | 0.942 |
| Transferases (EC 2) | 0.987 | 0.952 | 0.959 | 0.955 | 0.948 |
| Hydrolases (EC 3) | 0.988 | 0.942 | 0.923 | 0.933 | 0.926 |
| Lyases (EC 4) | 0.998 | 0.975 | 0.958 | 0.967 | 0.965 |
| Isomerases (EC 5) | 0.998 | 0.972 | 0.962 | 0.967 | 0.966 |
| Ligases (EC 6) | 0.999 | 0.988 | 0.985 | 0.987 | 0.986 |
| Translocases (EC 7) | 0.999 | 0.965 | 0.975 | 0.97 | 0.97 |
| Unclassified Enzyme | 0.978 | 0.704 | 0.608 | 0.653 | 0.643 |
| Non-enzyme | 0.975 | 0.979 | 0.978 | 0.978 | 0.95 |

**Table 9:** Per-class metrics for the Protein Category head.

| **Protein Category** | Acc | Prec | Rec | F<sub>1</sub> | MCC |
| --- | --- | --- | --- | --- | --- |
| Enzymes | 0.973 | 0.967 | 0.969 | 0.968 | 0.945 |
| Transporters | 0.991 | 0.947 | 0.91 | 0.928 | 0.923 |
| Receptors | 0.982 | 0.79 | 0.772 | 0.781 | 0.772 |
| Signaling Proteins | 0.933 | 0.688 | 0.707 | 0.697 | 0.66 |
| Structural Proteins | 0.958 | 0.923 | 0.845 | 0.882 | 0.858 |
| Virion | 0.997 | 0.884 | 0.829 | 0.856 | 0.855 |
| Motor Proteins | 0.988 | 0.768 | 0.647 | 0.703 | 0.699 |
| Regulatory Proteins | 0.932 | 0.875 | 0.849 | 0.862 | 0.817 |

**Table 10:** Per–GO–term metrics for the test set.

| GO Term (ID) | Acc | Prec | Rec | F<sub>1</sub> | MCC |
| --- | --- | --- | --- | --- | --- |
| Oxidoreductase activity | 0.992 | 0.792 | 0.758 | 0.775 | 0.771 |
| Kinase activity | 0.985 | 0.927 | 0.808 | 0.863 | 0.858 |
| Protein serine/threonine kinase activity | 0.994 | 0.699 | 0.82 | 0.755 | 0.755 |
| Protein tyrosine kinase activity | 0.999 | 0.667 | 0.608 | 0.636 | 0.636 |
| Phosphatase activity | 0.995 | 0.912 | 0.785 | 0.844 | 0.844 |
| Protein serine/threonine phosphatase activity | 0.999 | 0.509 | 0.703 | 0.591 | 0.598 |
| Protein tyrosine phosphatase activity | 0.999 | 0.725 | 0.798 | 0.76 | 0.761 |
| Protease (peptidase) activity | 0.992 | 0.882 | 0.795 | 0.836 | 0.833 |
| Serine-type endopeptidase activity | 0.999 | 0.924 | 0.925 | 0.924 | 0.924 |
| Cysteine-type endopeptidase activity | 0.998 | 0.642 | 0.557 | 0.596 | 0.597 |
| Aspartic-type endopeptidase activity | 1 | 0.98 | 0.851 | 0.911 | 0.913 |
| DNA-directed DNA polymerase activity | 0.999 | 0.895 | 0.801 | 0.845 | 0.846 |
| DNA-directed RNA polymerase activity | 0.98 | 0.792 | 0.717 | 0.753 | 0.744 |
| Topoisomerase activity | 1 | 0 | 0 | 0 | 0 |
| Helicase activity | 0.999 | 0.913 | 0.942 | 0.928 | 0.927 |
| GTPase activity | 0.997 | 0.939 | 0.88 | 0.908 | 0.907 |
| ATPase activity | 0.993 | 0.918 | 0.887 | 0.903 | 0.899 |
| Methyltransferase activity | 0.987 | 0.899 | 0.77 | 0.83 | 0.826 |
| Acetyltransferase activity | 0.964 | 0.505 | 0.526 | 0.515 | 0.496 |
| Ubiquitin-protein transferase activity | 0.992 | 0.607 | 0.623 | 0.615 | 0.611 |
| RNA-dependent RNA polymerase activity | 1 | 1 | 0.889 | 0.941 | 0.943 |
| ATP binding | 0.991 | 0.97 | 0.967 | 0.968 | 0.963 |
| GTP binding | 0.998 | 0.977 | 0.955 | 0.966 | 0.965 |
| DNA binding | 0.981 | 0.885 | 0.864 | 0.875 | 0.864 |
| RNA binding | 0.984 | 0.813 | 0.784 | 0.798 | 0.79 |
| Metal ion binding | 0.966 | 0.832 | 0.813 | 0.822 | 0.804 |
| Zinc ion binding | 0.992 | 0.918 | 0.859 | 0.888 | 0.884 |
| Iron ion binding | 0.998 | 0.954 | 0.925 | 0.939 | 0.939 |
| Heme binding | 0.999 | 0.968 | 0.92 | 0.944 | 0.944 |
| NAD(P) binding | 0.998 | 0.942 | 0.929 | 0.936 | 0.935 |
| Coenzyme binding | 1 | 0 | 0 | 0 | 0 |
| Transmembrane transporter activity | 0.993 | 0.862 | 0.84 | 0.851 | 0.847 |
| ABC-type transporter activity | 0.998 | 0.861 | 0.93 | 0.894 | 0.894 |
| Ion channel activity | 0.999 | 0.914 | 0.733 | 0.813 | 0.818 |
| Proton-transporting ATPase activity | 1 | 0.954 | 0.966 | 0.96 | 0.96 |
| Calcium-transporting ATPase activity | 1 | 0.854 | 0.837 | 0.845 | 0.845 |
| G protein-coupled receptor activity | 0.999 | 0.828 | 0.726 | 0.774 | 0.775 |
| Receptor tyrosine kinase activity | 1 | 0.577 | 0.672 | 0.621 | 0.623 |

Table 10 (continued)
| GO Term (ID) | Acc | Prec | Rec | F<sub>1</sub> | MCC |
| --- | --- | --- | --- | --- | --- |
| Hormone receptor activity | 1 | 0.808 | 0.538 | 0.646 | 0.659 |
| Cytokine receptor activity | 1 | 0.611 | 0.846 | 0.71 | 0.719 |
| Transcription factor activity, DNA binding | 0.991 | 0.779 | 0.849 | 0.812 | 0.808 |
| Signal transducer activity | 1 | 0 | 0 | 0 | 0 |
| Protein dimerization activity | 0.999 | 0.896 | 0.893 | 0.894 | 0.894 |
| Chaperone binding / unfolded protein binding | 0.998 | 0.969 | 0.868 | 0.916 | 0.917 |
| Heat shock protein binding | 0.999 | 0.857 | 0.593 | 0.701 | 0.712 |
| Disulfide oxidoreductase activity | 1 | 0.868 | 0.702 | 0.776 | 0.781 |
| Nitric-oxide synthase activity | 1 | 0.929 | 0.542 | 0.684 | 0.709 |
| Phospholipase activity | 1 | 0.639 | 0.535 | 0.582 | 0.584 |
| Glycosyltransferase activity | 0.999 | 0.707 | 0.572 | 0.632 | 0.635 |
| Glycosidase activity | 1 | 0.917 | 0.786 | 0.846 | 0.849 |
| Hydrolase activity, acting on carbon-nitrogen bonds | 1 | 0.538 | 0.483 | 0.509 | 0.51 |
| Aminotransferase activity | 1 | 0.979 | 0.94 | 0.96 | 0.96 |
| Decarboxylase activity | 0.999 | 0.948 | 0.933 | 0.941 | 0.94 |
| Vitamin binding | 1 | 0 | 0 | 0 | 0 |
| Flavin adenine dinucleotide binding | 0.999 | 0.921 | 0.915 | 0.918 | 0.917 |
| FMN binding | 1 | 0.975 | 0.95 | 0.962 | 0.962 |
| Biotin binding | 1 | 1 | 0.25 | 0.4 | 0.5 |
| Tetrapyrrole binding | 1 | 0 | 0 | 0 | 0 |
| Antioxidant activity | 1 | 0.615 | 0.381 | 0.471 | 0.484 |
| Structural Molecule Activity | 0.998 | 0.833 | 0.678 | 0.747 | 0.75 |
| Motor activity | 1 | 0.977 | 0.925 | 0.95 | 0.951 |
| Nuclease activity | 0.999 | 0.891 | 0.709 | 0.789 | 0.794 |
| Translation factor activity, nucleic acid binding | 1 | 0.429 | 0.25 | 0.316 | 0.327 |
| Ligase activity | 0.996 | 0.945 | 0.927 | 0.936 | 0.934 |
| Lyase activity | 0.997 | 0.896 | 0.802 | 0.846 | 0.846 |
| Hydrolase activity, acting on ester bonds | 1 | 0.781 | 0.714 | 0.746 | 0.747 |
| DNA ligase | 1 | 0.968 | 0.933 | 0.95 | 0.95 |
| Actin binding | 0.997 | 0.549 | 0.561 | 0.555 | 0.553 |
| Microtubule binding | 0.998 | 0.704 | 0.623 | 0.661 | 0.661 |
| Symporter activity | 0.999 | 0.847 | 0.776 | 0.809 | 0.81 |
| Antiporter activity | 0.999 | 0.871 | 0.794 | 0.831 | 0.831 |
| Voltage-gated ion channel activity | 1 | 0 | 0 | 0 | 0 |
| Protein binding | 0.981 | 0.771 | 0.534 | 0.631 | 0.633 |
| Ubiquitin ligase E3 activity | 0.996 | 0.632 | 0.636 | 0.634 | 0.632 |
| Peroxidase activity | 0.999 | 0.922 | 0.883 | 0.902 | 0.902 |
| Superoxide dismutase activity | 1 | 0.986 | 0.922 | 0.953 | 0.954 |
| Phospholipid binding | 0.999 | 0.874 | 0.606 | 0.716 | 0.727 |

Table 10 (continued)
| GO Term (ID) | Acc | Prec | Rec | F<sub>1</sub> | MCC |
| --- | --- | --- | --- | --- | --- |
| Lipid transfer protein activity | 1 | 0.75 | 0.818 | 0.783 | 0.783 |
| Transferase activity | 0.986 | 0.941 | 0.915 | 0.928 | 0.921 |
| mRNA binding | 0.992 | 0.722 | 0.669 | 0.695 | 0.691 |
| RNA helicase activity | 0.999 | 0.922 | 0.895 | 0.909 | 0.908 |
| Histone acetyltransferase activity | 0.999 | 0.49 | 0.439 | 0.463 | 0.463 |
| Histone deacetylase activity | 1 | 0.864 | 0.708 | 0.779 | 0.782 |
| Cell adhesion molecule binding | 0.999 | 0.404 | 0.319 | 0.357 | 0.359 |
| Caspase activity | 1 | 0.75 | 0.3 | 0.429 | 0.474 |
| SNARE binding | 0.999 | 0.723 | 0.767 | 0.744 | 0.744 |

**Table 11:** Per-pathway performance on the held-out test set.

| Pathway Membership | Acc | Prec | Rec | F<sub>1</sub> | MCC |
| --- | --- | --- | --- | --- | --- |
| Glycolysis / Gluconeogenesis | 0.998 | 0.946 | 0.878 | 0.91 | 0.91 |
| TCA (Citric Acid) Cycle | 0.998 | 0.938 | 0.879 | 0.908 | 0.907 |
| Pentose Phosphate Pathway | 0.999 | 0.965 | 0.939 | 0.952 | 0.952 |
| Oxidative Phosphorylation | 0.993 | 0.946 | 0.908 | 0.926 | 0.923 |
| Fatty Acid Biosynthesis | 0.997 | 0.928 | 0.789 | 0.853 | 0.854 |
| Fatty Acid Beta-Oxidation | 1 | 0.887 | 0.901 | 0.894 | 0.894 |
| Shikimate Pathway | 1 | 0.991 | 0.975 | 0.983 | 0.983 |
| Purine Metabolism | 0.993 | 0.939 | 0.912 | 0.925 | 0.922 |
| Pyrimidine Metabolism | 0.998 | 0.969 | 0.932 | 0.95 | 0.949 |
| Folate / One-Carbon Metabolism | 0.997 | 0.953 | 0.94 | 0.947 | 0.945 |
| Mevalonate Pathway | 0.999 | 0.735 | 0.76 | 0.747 | 0.747 |
| Non-mevalonate (MEP/-DOXP) Pathway | 1 | 0.982 | 0.985 | 0.983 | 0.983 |
| Peptidoglycan Biosynthesis | 0.998 | 0.967 | 0.919 | 0.943 | 0.942 |
| Chitin Biosynthesis | 0.998 | 0.945 | 0.818 | 0.877 | 0.878 |
| Ubiquinone and Terpenoid-Quinone Biosynthesis | 0.997 | 0.944 | 0.911 | 0.927 | 0.926 |
| Glyoxylate and Dicarboxylate Metabolism | 0.999 | 0.906 | 0.876 | 0.891 | 0.89 |
| Glycerophospholipid Metabolism | 0.995 | 0.868 | 0.806 | 0.836 | 0.834 |
| Sphingolipid Metabolism | 0.998 | 0.714 | 0.678 | 0.695 | 0.695 |
| Branched-Chain Amino Acid Biosynthesis | 0.997 | 0.948 | 0.844 | 0.893 | 0.893 |

Table 11 (continued)
| Pathway Membership | Acc | Prec | Rec | F<sub>1</sub> | MCC |
| --- | --- | --- | --- | --- | --- |
| Sulfur Amino Acid Biosynthesis | 0.993 | 0.962 | 0.908 | 0.934 | 0.931 |

#### AstraSUIT2 Results

**Table 12:** Overall performance of AstraSUIT2 task heads.

| Task | Acc | Prec | Rec | F<sub>1</sub> | MCC |
| --- | --- | --- | --- | --- | --- |
| Cofactor | 0.993 | 0.982 | 0.982 | 0.982 | 0.977 |
| Cofactor Specific | 0.999 | 0.95 | 0.921 | 0.936 | 0.935 |
| Domain | 0.981 | 0.866 | 0.869 | 0.868 | 0.858 |
| Host | 0.974 | 0.869 | 0.906 | 0.887 | 0.873 |
| Membrane Type | 0.989 | 0.965 | 0.968 | 0.967 | 0.96 |
| Subcellular Location | 0.984 | 0.882 | 0.864 | 0.873 | 0.864 |
| TM Helices | 0.994 | 0.877 | 0.878 | 0.878 | 0.875 |
| Quaternary Category | 0.973 | 0.886 | 0.856 | 0.871 | 0.856 |
| Quaternary Stoichiometry | 0.993 | 0.917 | 0.875 | 0.895 | 0.892 |

**Table 13:** Performance by cofactor type.

| Cofactor Class | Acc | F<sub>1</sub> | MCC |
| --- | --- | --- | --- |
| Organic | 0.997 | 0.961 | 0.959 |
| Metal Ions | 0.987 | 0.955 | 0.947 |
| Iron-Sulfur Clusters | 0.999 | 0.976 | 0.975 |
| Nucleotides | 0.999 | 0.904 | 0.904 |

**Table 14:** Performance by cofactor label.

| Cofactor | Acc | F<sub>1</sub> | MCC |
| --- | --- | --- | --- |
| Zinc | 0.995 | 0.932 | 0.93 |
| Magnesium | 0.992 | 0.938 | 0.933 |
| Calcium | 0.995 | 0.837 | 0.834 |
| Iron | 0.998 | 0.951 | 0.95 |
| Manganese | 0.995 | 0.894 | 0.893 |
| Copper | 0.999 | 0.854 | 0.855 |
| Nickel | 1 | 0.875 | 0.876 |
| Cobalt | 0.999 | 0.824 | 0.824 |

Table 14 (continued)
| **Cofactor** | Acc | F<sub>1</sub> | MCC |
| --- | --- | --- | --- |
| Molybdenum | 1 | 0.894 | 0.895 |
| Selenium | 1 | 0 | 0 |
| Tungsten | 1 | 0 | 0 |
| Vanadium | 0.999 | 0.919 | 0.919 |
| FAD | 0.999 | 0.969 | 0.968 |
| FMN | 1 | 0.937 | 0.937 |
| NAD | 0.999 | 0.974 | 0.973 |
| NADP | 0.999 | 0.925 | 0.924 |
| Heme | 0.999 | 0.951 | 0.951 |
| Heme B | 0.999 | 0.939 | 0.939 |
| Heme C | 1 | 0.866 | 0.866 |
| Biotin | 0.999 | 0.955 | 0.955 |
| Tetrahydrofolate | 0.999 | 0.946 | 0.946 |
| Coenzyme A | 1 | 0.821 | 0.822 |
| PLP | 1 | 0.95 | 0.95 |
| TPP | 1 | 0.84 | 0.841 |
| ThDP | 1 | 0.927 | 0.928 |
| SAM | 1 | 0.97 | 0.97 |
| Lipoic Acid | 1 | 0.87 | 0.87 |
| Molybdopterin | 1 | 0.667 | 0.707 |
| PQQ | 1 | 0.3 | 0.314 |
| Cobalamin | 1 | 0 | 0 |
| 4Fe-4S Cluster | 1 | 0.965 | 0.965 |
| 2Fe-2S Cluster | 1 | 0.971 | 0.971 |
| Fe-S Cluster | 1 | 0.908 | 0.908 |
| Others | 1 | 0.675 | 0.678 |
| ATP | 1 | 0.861 | 0.861 |
| ADP | 1 | 0.6 | 0.601 |
| AMP | 1 | 0.931 | 0.932 |
| GTP | 1 | 0.581 | 0.621 |
| GDP | 1 | 0.976 | 0.976 |
| GMP | 1 | 1 | 1 |
| CTP | 1 | 0 | 0 |
| CDP | 1 | 0 | 0 |
| CMP | 1 | 0.667 | 0.707 |
| UTP | 1 | 0.556 | 0.57 |
| UDP | 1 | 0.4 | 0.436 |
| UMP | 1 | 0 | 0 |
| TTP | 1 | 0 | 0 |
| dTTP | 0.982 | 0.946 | 0.936 |

**Table 15:** Performance by domain classification label.

| **Domain Label** | Acc | F<sub>1</sub> | MCC |
| --- | --- | --- | --- |
| Bacteria: Gram-negative | 0.967 | 0.938 | 0.915 |
| Bacteria: Gram-positive | 0.968 | 0.832 | 0.814 |
| Bacteria: Other | 0.941 | 0.759 | 0.726 |
| Eukarya: Mammals | 0.965 | 0.893 | 0.873 |
| Eukarya: Plants | 0.979 | 0.916 | 0.904 |
| Eukarya: Fungi | 0.983 | 0.894 | 0.885 |
| Eukarya: Parasites | 0.996 | 0.808 | 0.808 |
| Eukarya: Other | 0.952 | 0.732 | 0.706 |
| Archaea | 0.995 | 0.916 | 0.914 |
| Viruses: DNA Viruses | 0.996 | 0.74 | 0.738 |
| Viruses: RNA Viruses | 0.999 | 0.895 | 0.894 |
| Viruses: Retroviruses | 1 | 0.872 | 0.873 |
| Viruses: Other | 0.995 | 0.693 | 0.691 |
| Synthetic/Engineered | 1 | 0 | 0 |

**Table 16:** Organism association.

| **Organism Label** | Acc | F<sub>1</sub> | MCC |
| --- | --- | --- | --- |
| Animal Pathogen | 0.994 | 0.645 | 0.646 |
| Animal-associated | 0.877 | 0.889 | 0.751 |
| Human Pathogen | 0.999 | 0.504 | 0.522 |
| Human-associated | 0.913 | 0.531 | 0.506 |
| Insect Pathogen | 0.988 | 0.701 | 0.696 |
| Plant Pathogen | 0.999 | 0.52 | 0.534 |
| Bacterial Pathogen | 0.999 | 0.347 | 0.348 |
| Bacterial-associated | 0.972 | 0.971 | 0.944 |
| Environmental | 0.994 | 0.829 | 0.826 |
| Commensal/Symbiont | 1 | 0.679 | 0.685 |

**Table 17:** Membrane type.

| **Membrane Type** | Acc | F<sub>1</sub> | MCC |
| --- | --- | --- | --- |
| Soluble | 0.973 | 0.983 | 0.916 |
| Single-pass | 0.988 | 0.881 | 0.875 |
| Multi-pass | 0.994 | 0.966 | 0.963 |
| Lipid-anchored | 1 | 0 | 0 |
| GPI-anchored | 1 | 0 | 0 |

Table 17 (continued)
| **Membrane Type** | Acc | F<sub>1</sub> | MCC |
| --- | --- | --- | --- |
| Peripheral Membrane | 0.978 | 0.824 | 0.813 |

**Table 18:** Transmembrane helix number.

| **TM Number** | Acc | F<sub>1</sub> | MCC |
| --- | --- | --- | --- |
| 1 TM Helix | 0.989 | 0.882 | 0.876 |
| 2–3 TM Helices | 0.993 | 0.828 | 0.824 |
| 4–6 TM Helices | 0.994 | 0.887 | 0.883 |
| 7 TM Helices | 0.996 | 0.81 | 0.809 |
| 8–12 TM Helices | 0.996 | 0.925 | 0.923 |
| >12 TM Helices | 0.999 | 0.87 | 0.87 |

**Table 19:**
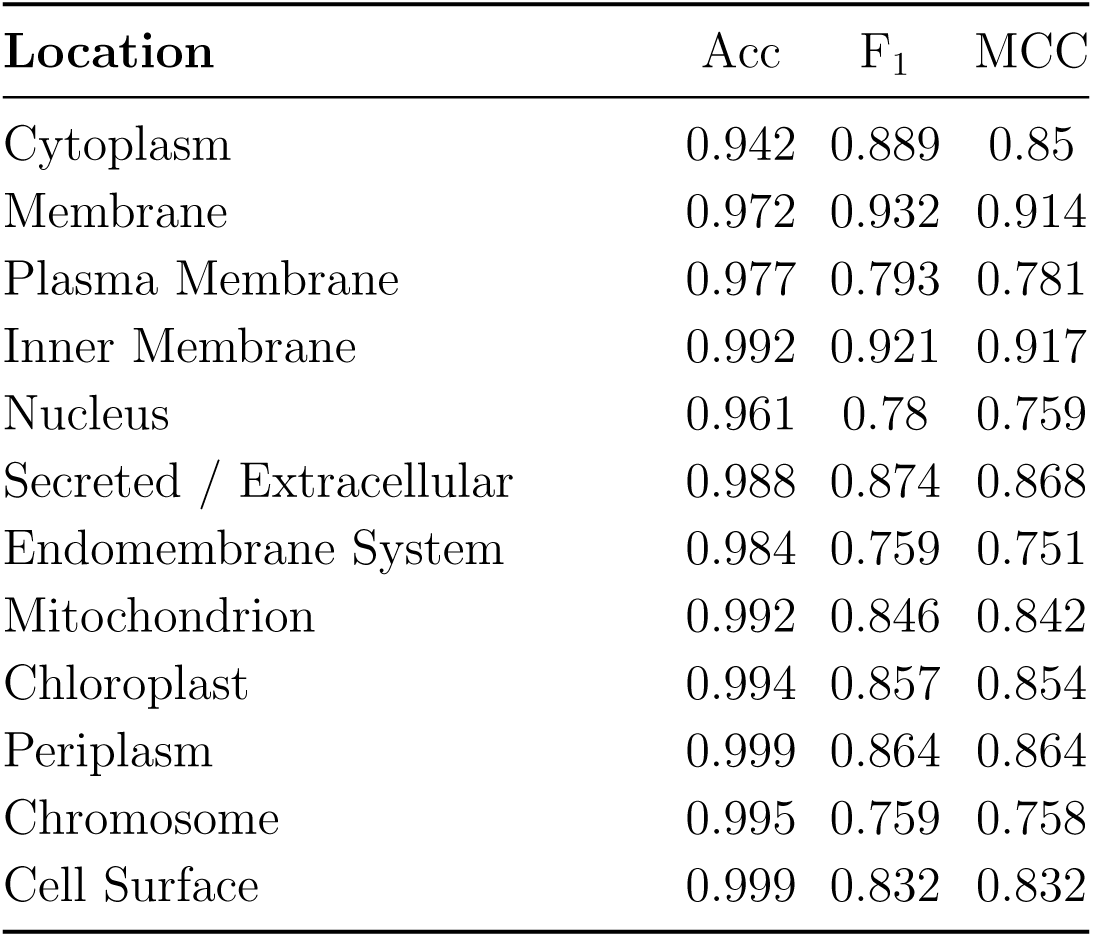
Subcellular localisation.

| **Location** | Acc | F<sub>1</sub> | MCC |
| --- | --- | --- | --- |
| Cytoplasm | 0.942 | 0.889 | 0.85 |
| Membrane | 0.972 | 0.932 | 0.914 |
| Plasma Membrane | 0.977 | 0.793 | 0.781 |
| Inner Membrane | 0.992 | 0.921 | 0.917 |
| Nucleus | 0.961 | 0.78 | 0.759 |
| Secreted / Extracellular | 0.988 | 0.874 | 0.868 |
| Endomembrane System | 0.984 | 0.759 | 0.751 |
| Mitochondrion | 0.992 | 0.846 | 0.842 |
| Chloroplast | 0.994 | 0.857 | 0.854 |
| Periplasm | 0.999 | 0.864 | 0.864 |
| Chromosome | 0.995 | 0.759 | 0.758 |
| Cell Surface | 0.999 | 0.832 | 0.832 |

**Table 20:** Quaternary structure type per label.

| **Quaternary type** | Acc | F<sub>1</sub> | MCC |
| --- | --- | --- | --- |
| Monomer | 0.992 | 0.898 | 0.894 |
| Homooligomer | 0.917 | 0.869 | 0.809 |
| Heretooligomer | 0.988 | 0.879 | 0.873 |
| Polymeric | 0.995 | 0.805 | 0.803 |

**Table 21:** Quaternary stoichiometry per label.

| **Stoichiometry** | Acc | F<sub>1</sub> | MCC |
| --- | --- | --- | --- |
| 1 | 0.992 | 0.898 | 0.895 |
| 2 | 0.975 | 0.893 | 0.879 |
| 3 | 0.997 | 0.877 | 0.875 |
| 4 | 0.993 | 0.913 | 0.91 |
| 5-6 | 0.997 | 0.922 | 0.921 |
| 7-12 | 0.998 | 0.929 | 0.928 |
| >12 | 1 | 0.965 | 0.965 |
| Indefinite | 0.995 | 0.805 | 0.802 |

